# Conformation and structural dynamics of the Xist lncRNA A-repeats

**DOI:** 10.1101/2022.07.26.501616

**Authors:** Alisha N. Jones, Frank Gabel, Stefan Bohn, Gregory Wolfe, Michael Sattler

## Abstract

LncRNAs are emerging to play crucial roles in the regulation of many essential cellular processes and have been linked to human disease, but a detailed understanding of their structure and how this relates to underlying molecular mechanisms is still limited. The structure that a lncRNA adopts can interconvert between multiple conformations. However, characterizing the structure and dynamics is challenging given their large size. Here, we present an integrated approach, combining biochemical and biophysical techniques to investigate the core structural elements and conformational dynamics of the A-repeats of the lncRNA Xist. We combine chemical RNA structure probing, SAXS, NMR-spectroscopy and cryo-EM to comprehensively describe the conformational landscape of the Xist A-repeats. We show that under native-like conditions, the A-repeats are modular, comprising building blocks made from stable AUCG tetraloop hairpins and inter-repeat dimers separated by flexible uracil-rich regions. The structural core of the A-repeats involves dimerization of sequential repeats to form two subdomains, comprising repeats 1-4 and 5-8. The overall topology of the A-repeats is dynamic, with structural variability linked to the uracil-rich linker regions. Our results rationalize context and buffer-dependent structural variations of the Xist lncRNA. The integrative approach presented here establishes a general pipeline for investigating lncRNA structure and dynamics.

**GRAPHICAL ABSTRACT:** 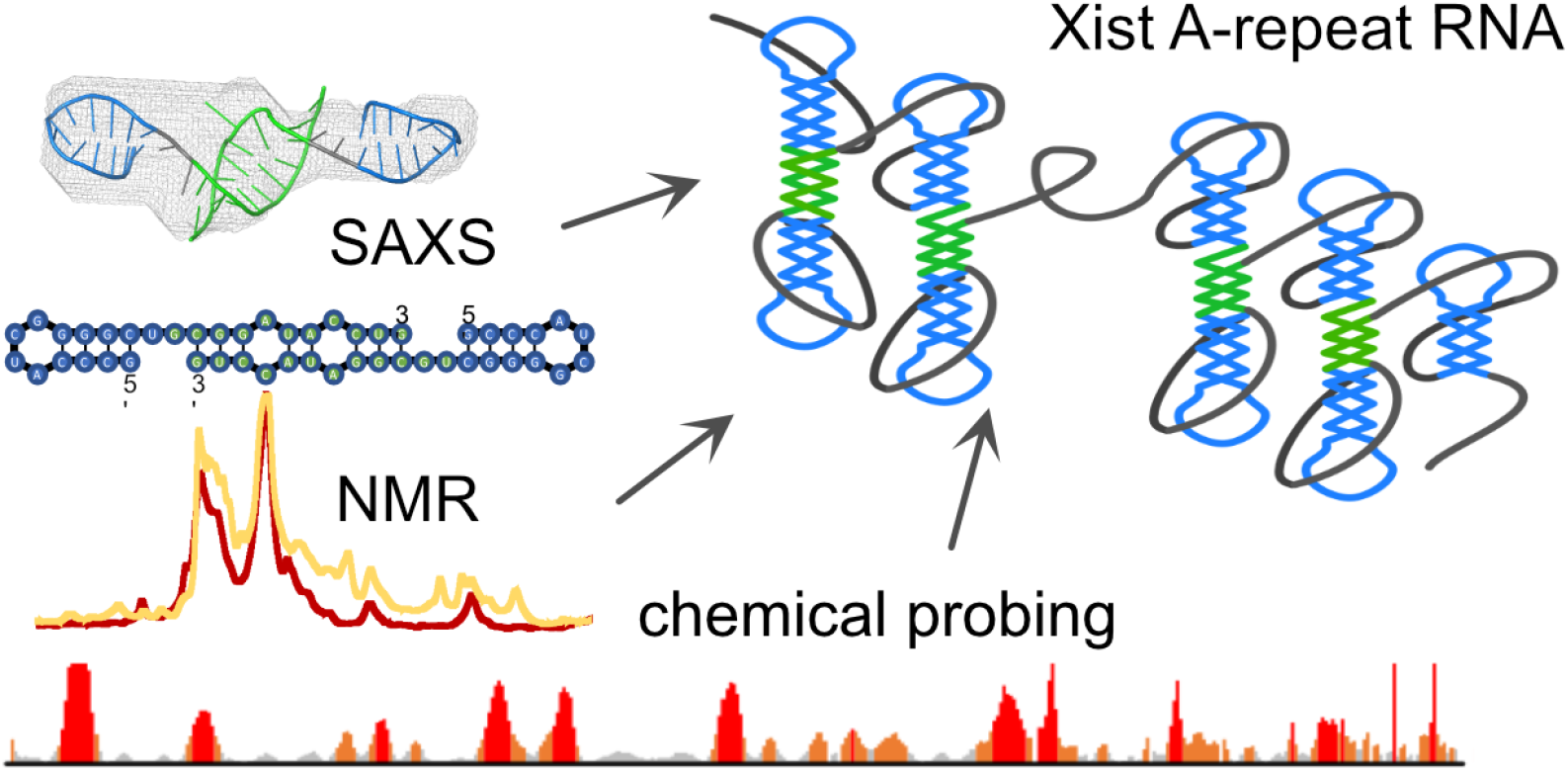

## INTRODUCTION

Long noncoding RNAs (lncRNAs) are non-protein-encoding transcripts with lengths of 500 or more nucleotides (nt), with some spanning up to several thousands of nucleotides ^1^. LncRNAs have emerged as important regulators of gene expression and cellular functions, as transcriptional regulators at the level of chromatin, as scaffolding platforms, and as guides for RNP localization ^2–9^. Recent advancements in chemical probing experiments and their coupling with next generation sequencing analysis have enabled us to determine the secondary structures of nearly 50 of the known 200,000 lncRNAs ^10–13^. Critically, the molecular functions of lncRNAs depend on local and global structures, which are modulated by RNA binding proteins ^14–22^.

RNAs are often conformationally flexible and may utilize structural variability to regulate biological function ^23–28^. For the lncRNA COOLAIR, it was recently demonstrated that changes in temperature promote the interconversion of structural states, with specific conformations being linked to specific biological processes ^29^. The lncRNA Braveheart is dynamic and interactions with its cognate protein binding partner, the cellular nucleic acid binding protein (CNBP), alter the conformation of the lncRNA ^17^. The structures of lncRNAs are also modulated by post-transcriptional modifications, such as N-6 methyladenosine (m^6^A), as has been shown for Xist and MALAT1 ^20,30^. In these instances, the m^6^A modification leads to conformational changes to increase the accessibility of protein binding sites. Thus, lncRNA conformational dynamics are critical for function.

In this work, we present a pipeline to identify core structural domains within a lncRNA, and to investigate the conformation landscape in the context of the full-length transcript. The presence of structural states that may be adopted during co-transcriptional folding is approximated by analyzing lncRNA fragments of different lengths. We demonstrate this approach with the human Xist A-repeats, a ∼ 450 nucleotide region that is located at the 5’ end of the Xist lncRNA. Xist is responsible for coating and transcriptionally silencing the inactive X-chromosome in placental mammals, a process known as XCI ^7,31–33^. In the absence of the A-repeats, the efficiency of silencing is drastically reduced. The A-repeats are composed of multiple 24-nucleotide conserved elements (7.5 elements in mouse and 8.5 elements in human) that are separated by uracil-rich sequences ^7^. The first 14 nucleotides of each A-repeat fold to form a stable AUCG tetraloop hairpin, while the second half is extended in solution to promote dimerization with other A-repeats ^34,35^. Notably, to date more than 15 secondary structure models have been proposed for the Xist A-repeats, which may correlate with conformational dynamics and variability, influenced by experimental conditions and approaches used ^34,36–42^.

We generated various stalled transcripts of the Xist A-repeat lncRNA to capture locally folded, structured elements that may fold cotranscriptionally. Combining selective 2’ hydroxyl acylation analyzed by primer extension (SHAPE) experiments ^43^, nuclear magnetic resonance (NMR), and small angle X-ray scattering (SAXS)) we identified locally folded, dimeric elements formed by two consecutive A-repeats. The dimeric units are comprised of the previously reported Xist A-repeat AUCG tetraloop structure, and are formed by dimerization via a 3’ region following the tetraloop fold. The conformational dynamics of the Xist A-repeat lncRNA are assessed by NMR, SAXS and cryogenic electron microscopy (cryo-EM). Based on these data, we propose that the Xist A-repeats sample multiple conformational states, which are expected to be modulated by the interaction with RNA binding proteins and base modifications play an important role for molecular mechanisms linked to X-inactivation.

## MATERIAL AND METHODS

### DNA template preparation

DNA encoding the full-length Xist transcript (human) was kindly provided by Dr Carolyn Brown. DNA templates corresponding to the full-length Xist A-repeats (nucleotides 330-796, accession number gi|340393|gb|M97168.1) and each of the A-repX RNAs were subcloned using primers purchased from Eurofins Genomics. The 5’ primer for generating full-length and A-repX RNAs was designed to contain the 17-nuclotide T7 promoter to facilitate *in vitro* transcription by the T7 polymerase. The 3’ primer for the transcripts analyzed by SHAPE encoded for a 3’ cassette (for primer annealing) following the final A-repeat. For A-repX transcripts analyzed by NMR and SAXS, this cassette was not present. High-fidelity Phusion polymerase (New England Biolabs) was used to amplify the DNA constructs according to manufacturer’s instructions. DNA was assessed and purified on a 1% TAE agarose gel, followed by extraction using the Wizard PCR DNA extraction and purification kit, according to manufacturer instructions.

### RNA preparation

RNAs were transcribed *in vitro* using in-house prepared T7 polymerase. Briefly, 800 ng of template DNA was supplemented with 20 mM MgCl_2_, 20X transcription buffer (100 mM Tris pH 8, 100 mM spermidine, 200 mM DTT), 5% PEG 8000, 10 mM of each rATP, rUTP, rGTP, and rCTP, and 0.03 mg of T7 polymerase. Transcriptions were incubated at 37°C for two hours followed by purification, NMR measurements, or the immediate addition of 1M7 as described below.

Full-length Xist RNAs analyzed using the buffer conditions listed in Table 1 were prepared as previously described and equilibrated in their respective buffers (see **Table 2**). Full-length Xist A-repeat RNA corresponding to the *ex vivo* Xist transcript analyzed by Smola et al., was purified under non-denaturing conditions using size exclusion chromatography, with buffer conditions matching the transcription reaction buffer listed above to maintain folding of the RNA (but excluding rNTPs and T7). RNAs were concentrated to 75 μM. RNAs were then immediately assessed by NMR or cryo-EM. SAXS samples were shipped at 4°C (to prevent degradation) for SAXS analysis.

**Table 1.** SAXS parameters obtained for each of the arrested A-rep RNAs studied.

| RNA construct (298 K) | $R_g$ (nm) |
| --- | --- |
| **14-mer** | $2.12 \pm 0.05$ |
| **24-mer** | $3.08 \pm 0.02$ |
| **A-rep1** | $4.10 \pm 0.07$ |
| **A-rep2** | $5.20 \pm 0.09$ |
| **A-rep3** | $6.11 \pm 0.19$ |
| **A-rep4** | $7.20 \pm 0.10$ |
| **A-rep5** | $7.86 \pm 0.20$ |
| **A-rep6** | $12.86 \pm 0.43$ |
| **A-rep8** | $11.39 \pm 0.13$ |

**Table 2.** Buffer conditions used to probe the secondary structure of the Xist A-repeats.

| Reference | Buffer condition |
| --- | --- |
| **Duszczek, et al.** | 298 K; 10 mM NaH<sub>2</sub>PO<sub>4</sub>/Na<sub>2</sub>HPO<sub>4</sub> pH 6.0, 100 mM NaCl, 0.02 mM EDTA, 0.02% azide |
| **Maenner, et al.** | 298 K; 20 mM HEPES-KOH pH 7.9, 100 mM KCl, 0.2 mM EDTA pH 8.0, 0.5 mM DTT, 0.5 mM PMSF, 20% glycerol, 3.25 mM MgCl<sub>2</sub> |
| **Liu, et al.** | 310 K/298 K; 25 mM K-HEPES pH 7.0, 0.1 mM Na-EDTA, 150 mM KCl, 15 mM MgCl<sub>2</sub> |
| **Smola, et al.** | 310 K; 100 mM HEPES pH 8.0, 100 mM NaCl, 10 mM MgCl<sub>2</sub> |

A-repX RNAs analyzed by SHAPE were treated immediately post-transcription (after two hours, when T7 activity was diminished) with an optimized concentration of 1M7 (25 mM) (or a DMSO control) for 5 minutes at 37°C. 25 mM 1M7 was selected to ensure single-hit modifications, to omit reactivity occurring due to cooperativity ^57^. Reactions were quenched with 6X denaturing buffer (8 M urea, 10 mM Tris pH 8.1, 40 mM EDTA), followed by purification using urea denaturing polyacrylamide gels (6.5 or 8%). Bands corresponding to the 1M7-modified RNA were excised from the gel and the RNA was extracted using the crush and soak method. Following extraction, RNAs were equilibrated against water and diluted to 1 μM.

### Selective 2’OH acylation analyzed by primer extension

1M7-treated A-repX RNAs were reverse transcribed with Superscript III reverse transcriptase (purchased from ThermoFisher) according to manufacturer’s instructions, using a 5’ FAM labeled primer (purchased from Eurofins genomics). Sequencing reactions (where the reverse transcription was carried out in the presence of ddATP, ddCTP, ddGPT, or ddTTP) were carried out to aid with nucleotide band assignments. The resulting cDNA was precipitated with ethanol and redissolved in 100% HiDi formamide for analysis by capillary electrophoresis. Capillary electrophoresis was carried out on an ABI 3130XL genetic analyzer. QuSHAPE ^58^ was used to obtain SHAPE reactivities. Scripts to generate folding restraint files used to fold each A-repX arrested RNA, as well as the SHAPE data, are available at github.com/gpwolfe/ctf.

### Nuclear magnetic resonance

All 1D NMR experiments were carried out on an 800 MHz spectrometer equipped with a cryogenic probe (Bruker) at 298 K and 280 K. 1D ^1^H spectra were recorded using a WATERGATE flip back sequence ^59^ on RNAs concentrated to 75 μM, and supplemented with 10% D_2_O.. Spectra were processed and analyzed using TopSpin (v3.7).

### Small angle X-ray scattering

Natively purified RNAs were concentrated to 0.5, 1.0, and 1.5 mg/mL in their respective buffers (see **Table 2**). Prior to measurement (carried out 298 K), RNAs were incubated at 310 K for 30 minutes. SAXS measurements were carried out at beamline P12 in Hamburg at the EMBL at the PETRA III storage ring. Buffer subtraction, scaling, and merging of data was performed in Primus.

### Cryo-Electron Microscopy

Natively purified RNAs (10 μM) were incubated in their respective buffers at 310 K for 30 min prior to grid preparation. For cryoEM sample preparation, 4.5μl of the buffered RNA solution was applied to glow discharged Quantifoil 2/1 grids, blotted for 3.5s with force 4 in a Vitrobot Mark III (Thermo Fisher) at 100% humidity and 4°C, and plunge frozen in liquid ethane, cooled by liquid nitrogen. Cryo-EM data was acquired with an FEI Titan Krios transmission electron microscope using SerialEM software ^60^. Movie frames were recorded at a nominal magnification of 22,500X using a K3 direct electro detector (Gatan). The total electron dose of ∼ 55 electrons per Å^2^ was distributed over 30 frames at a calibrated physical pixel size of 1.0 Å. Micrographs were recorded in a defocus range of - 0.5 to -3.0 μm. Particles were picked using Cryolo ^61^ and micrographs analyzed in Relion ^62^.

## RESULTS

### Stalled RNA transcripts reveal core structured elements formed by the A-repeat

We identified core structured elements formed by of the Xist A-repeats by evaluating the folding of secondary structure elements of stalled RNA fragments of various lengths using SHAPE chemical probing, NMR, and SAXS. For this, we prepared RNA transcripts that were stalled at various positions along the A-repeats (extending from the 5’ end of the RNA through the end of each A-repeat–Xist A- rep1 to Xist A-rep8) in order to gain insight into the hierarchical folding patterns adopted by the Xist A-repeat RNA. Immediately following *in vitro* transcription, RNA constructs were probed with 1- methyl-7-anhydride (1M7). Using this approach, omitting the need for denaturing purification, the RNA is expected to be probed in its native folding state.

Inspection of the SHAPE reactivity profiles for each stalled A-rep RNA shows that nucleotides of the uracil-rich region exhibit increased SHAPE reactivities relative to nucleotides that comprise the A-repeat elements. This indicates that the uracil-rich linkers are single stranded. Specific to the A-repeat elements, we find increased SHAPE reactivity for nucleotides that comprise the AUCG tetraloop of each major hairpin (**Figure 1A**, aligned with blue boxes). Residues corresponding to the second half of the repeat (**Figure 1A**, aligned with green boxes) show little to no SHAPE reactivity relative to the AUCG nucleotides. These data suggest that the A-repeats adopt a stable secondary structure, consistent with the previously identified stable AUCG stem loop structure formed by the major hairpin and the predicted minor hairpin being involved in secondary structure. Strikingly, a comparison of the SHAPE reactivity profiles of consecutive extended A-repeat RNA fragments reveals that the structured elements adopted by the Xist A-repeats are locally folded, as the SHAPE reactivity profile at the 5’ end of the RNA does not change when additional nucleotides are appended to the 3’ end of the transcript. The 3’ ends of each RNA show increased SHAPE reactivity due to intrinsic flexibility, thereby indicating a lack of base-paired nucleotides. Upon extension of the transcript, new structured domains are formed among the appended nucleotides, whenever an even number of repeats is included (**Figure 1A**).

**Figure 1.**
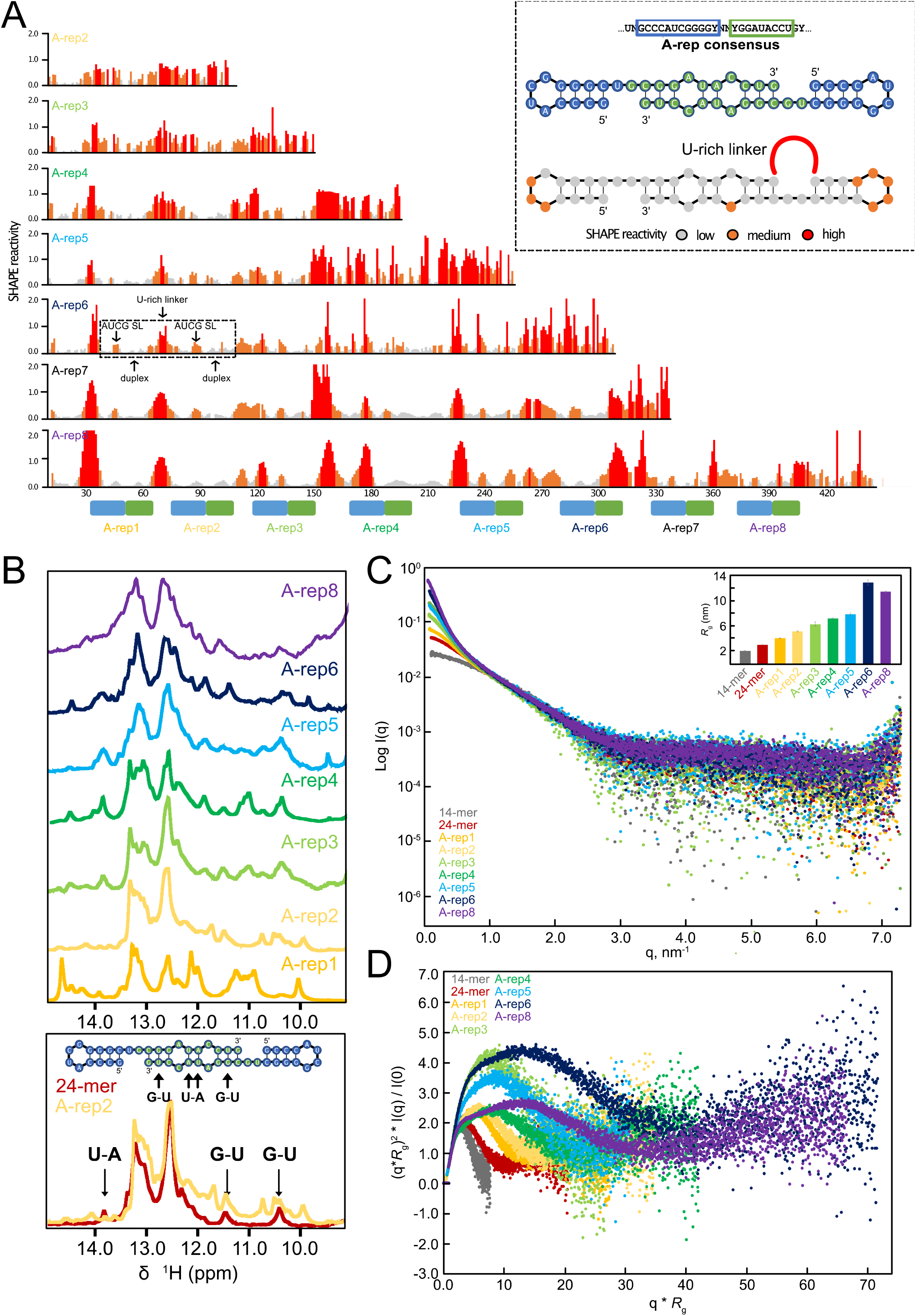
SHAPE and SAXS analysis of stalled fragments of the Xist A-repeats. **(A)** The SHAPE reactivity profile for extended Xist A-repeat RNA fragments. The inset box shows the SHAPE reactivity profile mapped to a duplex formed by two consecutive A-repeats. **(B)** ^1^H 1D NMR imino fingerprint spectra for each of the Xist A-repeat RNAs. **(C)** SAXS curves for extended Xist A-repeat RNA fragments. Inset bar graph shows *R*_*g*_ values for each fragment. **(D)** Dimensionless Kratky plot for extended Xist A-repeat RNA fragments.

It is noteworthy that the SHAPE reactivity profiles for the eight 24-nucleotide A-repeats are highly similar, with the linkers between them showing increased SHAPE reactivity. We previously reported that the 24-nucleotide A-repeats dimerize with subsequent A-repeats via the 3’ region of each repeat ^35^, confirmed by imino NMR spectra that show base pairing for the 3’ region. We thus collected 1D imino ^1^H NMR experiments for each of the extended A-rep transcripts to analyze and compare the base pairing in the context of these larger regions (**Figure 1B**). Indeed, the imino NMR signal fingerprint observed for the duplex formed by a single A-repeat dimer is also observed when multiple dimeric repeat units are present as the length of the RNA increases. For example, imino signals of an RNA comprising two repeats are highly similar and show the presence of U-A base pairs in the duplex regions formed by dimerization of the 3’ region of the repeats (**Figure 1B**, bottom). This is also seen for larger RNA constructs, comprising 3-8 repeats, although spectral quality is challenged by line-broadening due to the increasing molecular weight of the larger constructs. The NMR data further support the SHAPE analysis and confirm that the A-repeats fold to form repetitive structures of AUCG tetraloop hairpins that involve dimerization with subsequent A-repeat elements.

We next used SAXS to investigate the overall conformation of each stalled fragment of the Xist A-repeat RNA (**Figure 1C, Supplementary Table 1**). As a control, we carried out SAXS on the 14-mer AUCG tetraloop hairpin previously determined by NMR ^34^. CRYSOL analysis ^44^ reveals a χ^2^ value of 1.24 for the NMR structure (PDB ID 2y95 ^34^), indicating excellent agreement between the structure and the SAXS data (**Supplementary Figure 1**). We then analyzed SAXS data for the various RNA transcripts. Superposition of the SAXS curves of the different RNAs reveals an expected increase in *I*(q) with increasing length of the RNA (**Figure 1C**). Along the same lines, a steady increase in the *R*_g_ is found as the length of the RNA increases (**Figure 1C**, inset, **Table 1**). Comparison of the dimensionless Kratky plot suggests a compact, globular conformation for the 14-mer major hairpin, the 24-mer duplex-forming A-repeat, and A-reps 1 and 2 (**Figure 1D**). However, as the length of the RNA increases through A-rep3, the Kratky plot shows characteristics of molecule with significant extended and flexible conformations present. This is indicated by the steady increase of the curve as q\**R*_g_ increases. This trend is also observed for A-rep5 and A-rep6. Notably, these RNA constructs also show high SHAPE reactivity at the 3’ end of the RNA (**Figure 1A**), indicating the presence of single-stranded and flexible nucleotides. On the other hand, A-rep4 and A-rep8 have similar dimensionless Kratky profiles, where the curves come down at the highest *R*_g_ values, indicating the RNAs are rather compact. Consistent with this, the SHAPE reactivity at the 3’ end of these RNA is relatively lower than what is observed for A-rep5 and A-rep6. These data suggest that the first 4 and the full-length A-repeats may fold to form a compact structure, whereas intermediates transcripts are more flexible, likely due to poor folding of the 3’ ends of the RNAs. This is consistent with the presence of two globular domains formed by 4 repeats that assemble in the full A-repeat RNA structure to form an overall globular fold comprised of repetitive inter-repeat dimers.

### A structural model for the folded Xist A-repeat RNA structure

The core structured elements that contribute to the folding of the A-repeats can be derived considering that nucleotides located at the 5’ end of the RNA remain folded despite the addition of nucleotides to the 3’ end of the transcript. The Pearson *R* value of the reactivity profiles of consecutive constructs shows good agreement, with poorer correlations observed when the 3’ end of the RNA is largely unstructured (**Supplementary Table 2**). To this end, we developed a pipeline to generate folding restraints that build upon the structure adopted by the Xist A-repeats upon the addition of 3’ nucleotides. These data were then used to fold each of the Xist A-rep arrested RNAs (**Supplementary Figure 2**). Briefly, the Xist A-rep2 RNA secondary structure was predicted using RNAstructure based on SHAPE and NMR data ^45^; SHAPE reactivity data were included as soft folding restraints and unambiguous base pairs identified by NMR were set as hard folding restraints, in particular for the AUCG tetraloop hairpin and the dimerization region. To predict the structure of the Xist A-rep3 RNA, the SHAPE reactivity profiles of Xist A-rep3 and A-rep2 RNA were compared using a sliding window approach; for stretches of at least four nucleotides that show comparable SHAPE reactivities between the two RNAs, their corresponding structured elements (i.e., base pairs that are maintained) were added as restraints in the folding restraint file for A-rep3. For nucleotides of Xist A- rep3 that showed variable SHAPE reactivities relative to Xist A-rep2, SHAPE data) used as soft restraints to guide folding of that region of the RNA. The Xist A-rep3 RNA structure was then modelled using the curated folding restraint file, and this process was iterated through the 8 arrested RNA Xist A-rep constructs.

**Figure 2.**
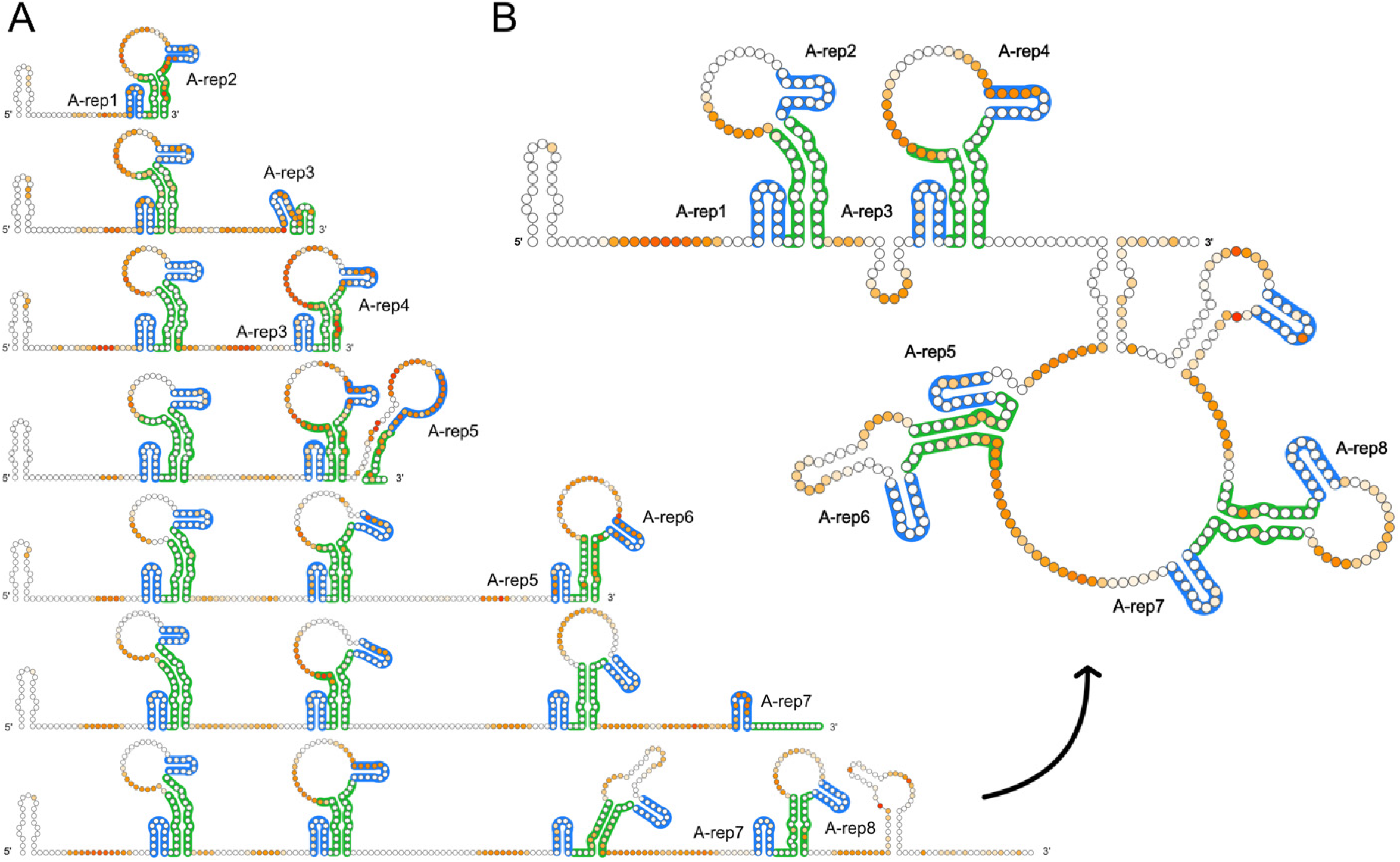
Folding of the Xist A-repeats. **(A)** The secondary structures of the Xist A-rep1 – A-rep8 RNA arrested RNAs; for simplicity, nucleotides between each A-rep duplex are represented as single stranded nucleotides. **(B)** Secondary structure of the full-length Xist A-repeats based on SHAPE restraints. Blue traces correspond to the 14-nt major hairpin, whereas green traces correspond to nucleotides involved in duplex formation.

The results of this analysis provide unique insight into the folding pathway of the Xist A-repeats as illustrated in **Figure 2**. We find that A-repeats 1 and 2, 3 and 4, 5 and 6, and 7 and 8 form duplex structures. As expected, nucleotides of the AUCG tetraloop hairpin show low SHAPE reactivity, consistent with our previous NMR structure which shows that these nucleotides are partially stacked and are less accessible to the SHAPE reagent ^34^. Interestingly, nearly all of the single-stranded uracil rich regions show reduced SHAPE reactivity. It has been recently shown that short stretches of uracil bases can stack under high salt concentrations (i.e., 200 mM), which may sterically prevent the accessibility of SHAPE reagents ^46^. These uracil bases may also form intra or inter U-U base pairs.

The integrated analysis of the SHAPE and NMR data suggests that the Xist A-repeats form a duplex structure, where the major hairpin of the first A-repeat is stably formed and coaxially stacked through the duplex formed by the region downstream of the tetraloop sequence with the hairpin from the sequential A-repeat ^34^. We used RNAComposer ^47^ to generate a tertiary structural model of two A-repeats forming a coaxially stacked duplex. The agreement of the RNAComposer-generated structural model with the experimental SAXS curve obtained for the 24-mer shows a χ^2^ = 9.3 (**Figure 3A**), while a monomeric 24-mer (structure generated in RNAComposer) shows a χ^2^ = 41.8 (**Figure 3B**). The relative comparison of the agreement with the experimental SAXS data supports the duplex existing as a folding unit. The rather larger χ^2^ value (9.26) likely reflects a poor prediction of structural details of the dimeric repeat by RNAComposer. DAMMIF *ab initio* bead modeling revealed a shape that strongly supports the coaxially stacked duplex (**Figure 3C**). Based on these data, alongside our NMR and SHAPE analysis, we propose the Xist A-repeat RNA folds by coaxial stacking of dimeric repeats, each comprising two AUCG tetraloop hairpins, and that the Xist A-repeat RNA overall adopts a modular structural arrangement, where two larger domains made of four repeats each form a compact higher order structure (**Figure 3D**).

**Figure 3.**
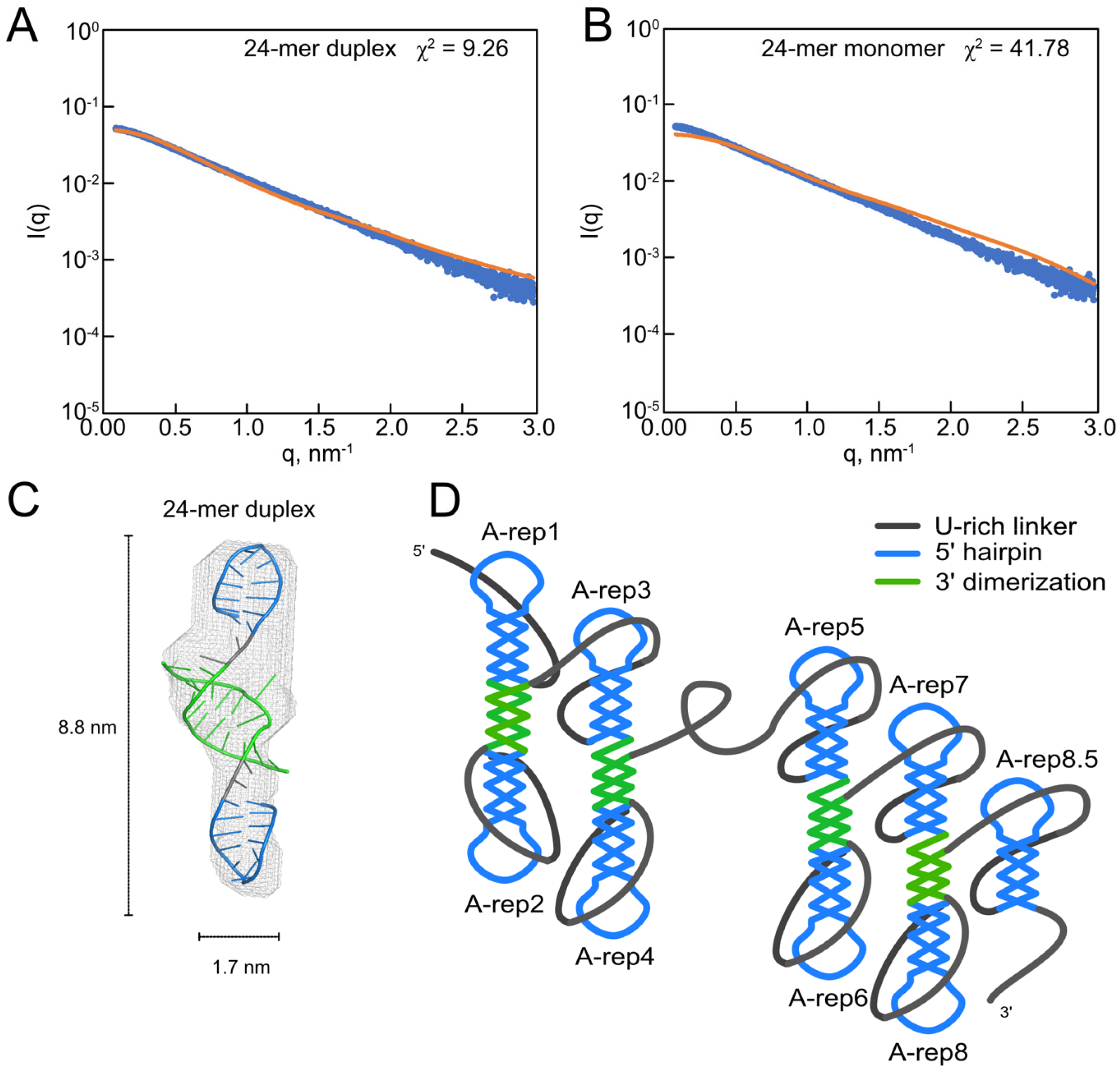
Structural model of the Xist A-repeats. **(A)** CRYSOL goodness of fit curve for the 24-mer SAXS curve (highest concentration curve at 1.98 mg / mL) with a duplexed 24-mer RNA structure and a **(B)** monomeric 24- mer structure. **(C)** The RNAComposer model for the duplexed 24-nucleotide Xist A-repeat RNA element fits well inside the 3D DAMMIF reconstruction of the structure based on SAXS data. **(D)** Cartoon of the proposed tertiary structure of the full-length Xist A-repeats.

### Xist A-repeat RNA structure is influenced by buffer composition *in vitro*

Inspection of the secondary structures predicted from our approach revealed some regions that do not match their SHAPE reactivity (i.e., single-stranded nucleotides showing little to no reactivity). This has been suggested to indicate structurally dynamic regions of RNA ^48,49^. Inspection of previously proposed secondary structures for Xist in cells and *in vitro* reveal either modular or non-modular arrangements of the A-repeats ^42^. Considering the reported structures vary significantly, we speculated whether the buffer conditions, under which the RNA was studied, influence the structure and dynamics of the Xist A-repeats (**Table 2**).

The secondary structure of the Xist A-repeats has been investigated *in vitro* by both chemical probing and enzymatic cleavage under four different buffer conditions, each resulting in a different secondary structure prediction ^34,36,37,39^ (**Table 2**). The most distinct differences between these buffer conditions concern the pH (ranging from 6.0 – 8.0) and salt content (100 to 150 mM KCl or NaCl, while MgCl_2_ is absent or ranges from 3.25 mM to 15 mM), two factors which are known to influence how an RNA folds ^39,50,51^. We therefore prepared the human Xist A-repeat RNA under the conditions described in each of the four studies, and used SAXS, NMR, and cryo-EM to study the effect of buffer conditions on the overall conformation of the RNA. Hereafter, these four conditions are referred to by the last name of the first author of each study: Maenner et al., Liu et al., Duszczyk et al., and Smola et al.

We first used SAXS to investigate the global conformation, flexibility and degree of folding of the Xist A-repeats under each of the four conditions at 298 K. Scattering profiles were recorded at three RNA concentrations between 0.20 and 2.1 mg/mL (**Supplementary Table 3**). The concentration series showed no inter-particle effects, and thus the highest concentration data were used for analysis (**Supplementary Figure 3A**,**B**). Guinier analysis reveals that the Xist A-repeat radius of gyration (*R*_g_), which is a measure of a molecule’s compactness, varies significantly depending on buffer composition (**Figure 4A, Table 2**). The Xist A-repeat RNA is most compact under the buffer conditions used by Duszczyk et al., with an *R*_g_ value of 7.56 ± 0.02 nm (**Table 3**), and increases for the Maenner et al., Smola et al., and Liu et al. buffer conditions with *R*_g_ values of 8.89 ± 0.18 nm, 9.83 ± 0.09 nm, and 12.10 ± 0.21 nm respectively, indicating a reduced compactness of the overall conformational ensemble in solution (**Table 3**). It was previously shown that the compactness of the RepA element of Xist (which is comprised of both the A and F repeat regions), as measured through the hydrodynamic radius (*R*_H_), decreases with an increase in MgCl_2_ concentration ^39^. By these standards, one might expect a similar trend to be observed for the four buffer conditions assayed in our SAXS studies: the Duszczyk buffer possesses no MgCl_2_, and the concentration of MgCl_2_ in the Maenner, Smola, and Liu buffers is 3.25 mM, 10 mM, and 15 mM, respectively. The *R*_*g*_ values we obtained show an opposite trend, and do not reflect a similar corollary relationship. However, buffer components and experimental conditions differ, and may thus affect the conformation in solution.

**Table 3.** SAXS parameters obtained for each of the RNAs studied.

| Buffer condition (298 K) | $R_g$ (nm) | $D_{max}$ (nm) |
| --- | --- | --- |
| **Liu et al.,** | 12.10 ± 0.21 | 42.98 |
| **Duszczek et al.,** | 7.56 ± 0.02 | 30.33 |
| **Smola et al.,** | 9.83 ± 0.09 | 33.03 |
| **Maenner et al.,** | 8.89 ± 0.18 | 34.69 |

**Table 4.** Buffer conditions and binding affinities for the Xist A-repeats with PRC2.

| Reference | Binding affinity | Buffer condition |
| --- | --- | --- |
| **Davidovich, et al.** | 9.1 nM | 50 mM Tris HCl, pH 7.5 (@ RT), 100 mM KCl, 2.5 mM MgCl<sub>2</sub>, 0.1 mM ZnCl<sub>2</sub>, 2 mM 2- mercaptoethanol, 0.1 mg/ml BSA, 0.1 mg/ml yeast tRNA, 0.05 % v/v NP40 and 5 % v/v glycerol |
| **Cifuentes-Rojas, et al.** | 80.9 nM | 50 mM Tris-HCl pH 8.0, 100 mM NaCl, 5 mM MgCl<sub>2</sub>, 10 µg/ml BSA, 0.05% NP40, 1 mM DTT, 20U RNaseOUT and 5% Glycerol in 20 µl |

**Figure 4.**
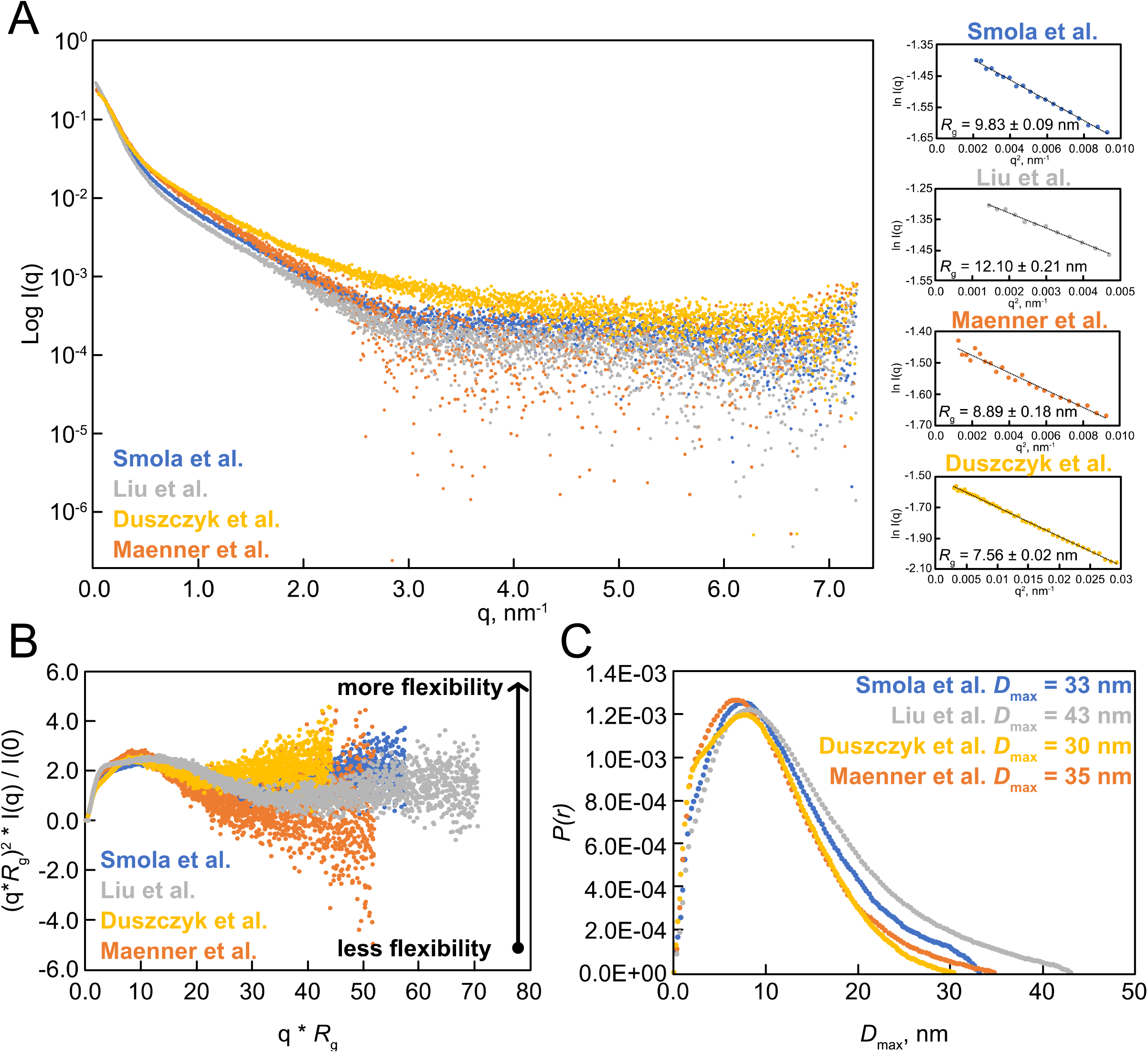
SAXS analysis of the Xist A-repeats at various buffer conditions at 298 K. **(A)** SAXS curves for buffers corresponding to Duszczyk et al., Maenner et al., Smola et al., and Liu et al. The radius of gyration, as determined from Guinier analysis, is shown on the right. The straight line is the linear regression fit in the range of q < 1.3/*R*_g_. **(B)** Dimensionless Kratky plot for each of the four buffered conditions. **(C)** *P(r)* curve for each of the four buffered conditions, with *D*_max_ values listed.

To assess the intrinsic flexibility of the Xist A-repeat RNA as a function of buffer condition, we generated dimensionless Kratky plots (**Figure 4B**) ^52^. For each of the buffer conditions, the dimensionless Kratky plot shows a profile characteristic of a partially compact particle with extended flexible elements. This is indicated by the steady increase of the curve as q\**R*_g_ increases through q\**R*_g_ = 10, followed by a plateau after a partial decrease at intermediate values. This result is not surprising, as lncRNAs are expected to be conformationally flexible ^17^. Interestingly, the dimensionless Kratky plots for each of the buffer conditions are not congruent, indicating significant structural differences depending on buffer conditions.

We investigated the degree of folding of the Xist A-repeats in each buffer by assessing the maximum distance (*D*_max_) from the *P(r)* pairwise distance distribution function (**Figure 4C**). The *D*_max_ is the maximum pairwise atom distance within the RNA. Comparable *D*_max_ values are observed for the conditions used by Maenner et al., Smola et al., and Duszczyk et al., reflecting maximum extended lengths of 35, 33, and 30 nm, respectively. The *D*_max_ length for the Liu et al. buffer condition is 43 nm, suggesting the Xist A-repeat RNA is more loosely structured. Altogether, these results support the notion that the Xist A-repeats, regardless of buffer composition, possess a high fraction of structured domains. However, the specific structural features seem to be significantly influenced by buffer conditions.

We next used single particle cryo-EM to obtain a complementary analysis of the RNA dimensions in the various buffer conditions. Observation and analysis of individual Xist A-repeat molecules in vitreous ice reveals that the RNA adopts a partially elongated structure with several branched domains originating from the main helical branch (**Figure 5B**). Analysis of the lengths of the helical stems and branches from ∼ 6,500 particles indicates a length distribution of 5 to 40 nm for all buffered conditions (**Figure 5B**). The particle length distribution is in excellent agreement with the *P(r)* distribution curves obtained from SAXS analysis. A sub-analysis of the longest chain for a subset of approximately 100 particles reveals that the Liu et al. buffered condition results in the most extended RNA particles, consistent with *D*_max_ values obtained from SAXS (**Figure 5B**, inset).

**Figure 5.**
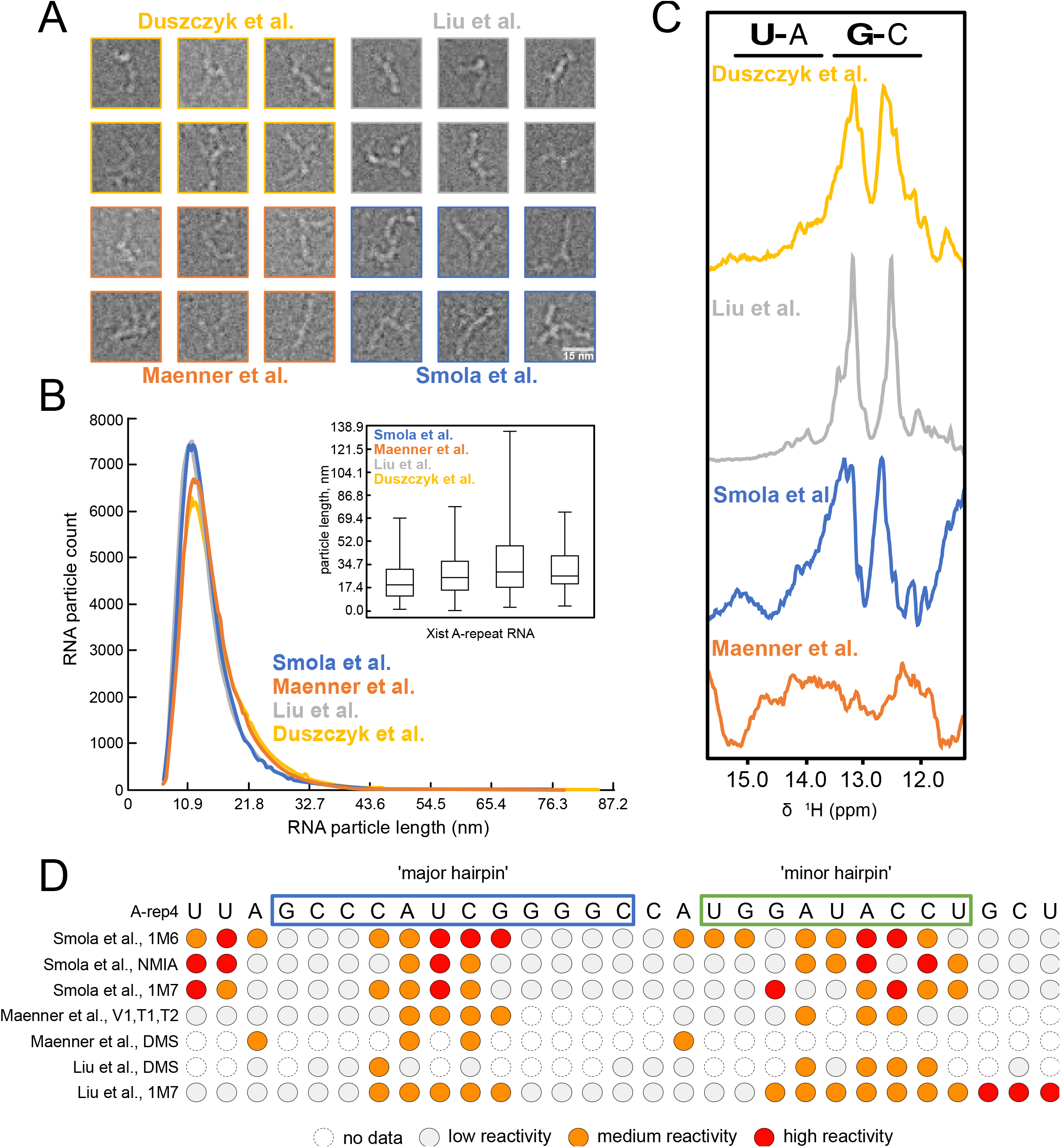
Effect of buffer on the structure of the Xist A-repeat RNAs. **(A)** Individual particles of the Xist A-repeats in vitreous ice, in buffers used by Maenner (orange), Smola (blue), Liu (grey), and Duszczyk (yellow) et al. **(B)** Distribution of maximum distances found in particles analyzed by cryo-EM. The inset shows the lengths of only the longest particles for approximately 100 particles for each buffered condition. **(C)** ^1^H 1D imino fingerprint spectra of the Xist A-repeats under varying buffer conditions at 298K. **(D)** Representative chemical probing (1M6, NMIA, 1M7, and DMS) and enzymatic cleavage (V1, T1, and T2) data for Xist A-repeat 4 as determined by Smola, et al., Maenner et al., and Liu et al. See also Data S2.

As analysis of SAXS data suggests the overall RNA conformational ensemble may vary depending on the buffer condition, we used ^1^H NMR to assess the presence of local RNA structure through observation of the imino proton resonances (**Figure 5C**). Imino protons of guanosine (H1) and uracil (H3) bases that are involved in base pair interactions are observable with chemical shifts between 10-15 ppm. The imino resonance profiles of the Xist A-repeats are not starkly different for the buffer conditions utilized by Smola et al., Liu et al., and Duszczyk et al., (note, that the glycerol present in the buffer used by Maenner et al. does not allow acquisition of high-quality NMR spectra). Two clusters of imino signals between 12.0-13.5 ppm are observed for Smola et al., Duszczyk et al., and Liu et al. conditions. Only minor differences are seen for the imino signals between each buffer, and the overall similarities are not surprising, as the chemical probing and enzymatic cleavage reactivity profiles reported by Smola et al., Liu et al., and Maenner et al. are also relatively similar (**Figure 5D, Data S1**). Taken together, the overall similarity of the imino NMR signal footprints indicates the presence of similar base pairing and folded RNA regions. However, the SAXS analyses reveal that the Xist A-repeats exhibit variable compactness under the different buffer conditions, suggesting variation in conformational ensembles and arrangements of individual local structural elements.

## DISCUSSION

The lncRNA Xist is one of the most well-studied long non-coding RNAs to date. Considering the critical role that the A-repeats have in transcriptional silencing, determining the structure and dynamic conformations of the A-repeats is important to further understand their interactions with RNA binding proteins and molecular mechanisms of XCI. Here, we consider the folding of local structural elements that may occur during transcription and employ an integrative approach to investigate the structure and dynamics of the Xist A-repeats. We show that the A-repeat RNA folds into two larger subdomains, each comprised of 4-5 repeats, with a modular arrangement of individual repeats by inter-repeat dimerization. Notably, the conformation and overall fold of the A-repeat is dynamic as indicated by our SAXS analysis, likely mediated by flexible U-rich linkers connecting the repeats. Overall, the similarity of chemical probing profiles *in vitro* and in cells for the mouse and human Xist A-repeats suggests that the basic topology and modular fold of the A-repeat RNA identified here is expected to represent the native structure *in vivo*.

Our SAXS, NMR, and cryo-EM data show that the conformational landscape of the Xist A-repeats varies under differential buffer conditions. For RNAs studied *in vitro*, purification of the transcript under denaturing conditions requires refolding of the RNA, which can lead to mis-folding, especially for longer RNAs ^53^. Even when purifying the RNA under native conditions, buffer composition (i.e., salt concentration, pH) and temperature can impact the structure adopted by the RNA. While in-cell chemical probing of the RNA secondary structure occurs under native conditions, RNAs can adopt multiple folded states at any given time in the cell, leading to structural information that reflects an average of structural states ^54,55^. The secondary structural data obtained for the Xist A-repeats, having been studied both *in vitro* and in cells, are subject to the limitations of each of these approaches.

Tertiary contacts and dimerization of individual structural elements of the A-repeat RNA provide an overall globular fold that nevertheless exhibits significant conformational flexibility. In addition to being modulated by protein binding events ^33^, the conformational ensemble of the Xist A-repeat RNA can be further finetuned by m^6^A modification to regulate the function of the lncRNA. It has recently been shown that the Xist A-repeats are subject to m^6^A modifications and that this has impacts on the structure adopted by the RNA ^20,56^, resulting in partial opening of the tetraloop structure formed by the A-repeats. The large differences we observe in the overall structure of the Xist A-repeats under varied buffer conditions further support the notion that the Xist A-repeats are highly dynamic and that their conformation can thus be modulated by protein binding or posttranscriptional modifications.

We propose that the integrative approach presented here provides an efficient way to study the conformational landscape of lncRNAs in solution. Combining chemical probing, NMR, and SAXS data enables the identification of core structured elements within lncRNA transcripts. The dynamics of these locally structured regions can be further evaluated using NGS read deconvolution algorithms ^29,49,54,55^. NMR, SAXS, and cryo-EM (even in the absence of 3D classifications) can be used to study the overall shape and dynamics of a lncRNA structural ensemble. The overall approach provides a foundation for linking conformational states of a lncRNA with molecular interactions and eventually biological activity.

## DATA AVAILABILITY

SHAPE reactivities and restraints files used to fold the Xist A-repeats are available at github.com/gpwolfe/ctf.

Experimental SAXS data have been deposited at SASDB under accession codes: SASDPB5, SASDPC5, SASDPD5, SASDPE5, SASDPF5, SASDPG5, SASDPH5, SASDPJ5, SASDPK5, SASDPL5, SASDPM5, SASDPN5, SASDPP5.

## SUPPLEMENTARY DATA

Supplementary Data are available at NAR online.

## AUTHOR CONTRIBUTIONS

Alisha Jones: Conceptualization, Formal analysis, Methodology, Validation, Visualization, Writing – original draft, review & editing. Frank Gabel: Formal analysis, Methodology, Validation. Stefan Bohn: Formal analysis, Methodology Validation. Gregory Wolfe: Formal analysis, Visualization. Michael Sattler: Conceptualization, Methodology, Validation, Writing – review and editing

## ACKNOWLEDGEMENTS

We thank Dr Carolyn Brown for providing the Xist DNA template, Dr Dominik Lenhart for the synthesis of 1M7 and Dr Arie Geerlof for preparation of T7 polymerase. We thank Johanna Voigt, Manuel Rohl and Markus Mueller for assistance with sample preparation. We thank Dr Gerd Gemmecker and Dr Sam Asami for help with NMR experiments, Dr Ralf Stehle for assistance with SAXS analysis, and members of the Sattler group for helpful discussions. The synchrotron SAXS data were collected at beamline P12 operated by EMBL Hamburg at the PETRA III storage ring (DESY, Hamburg, Germany). We would like to thank Stefano Da Vela for the assistance in using the beamline. We acknowledge access to NMR spectrometers at the Bavarian NMR center, and to cryo-EM instrumentation at the MPI for biochemistry, Martinsried.

