## Supplementary Information for "Conformation and structural dynamics of the Xist lncRNA A-repeats"

**This PDF file includes:**

Supplementary Figs. S1 to S3
Supplementary Tables S1 to S3

**Other Supplementary Materials for this manuscript include the following:**
Data S1

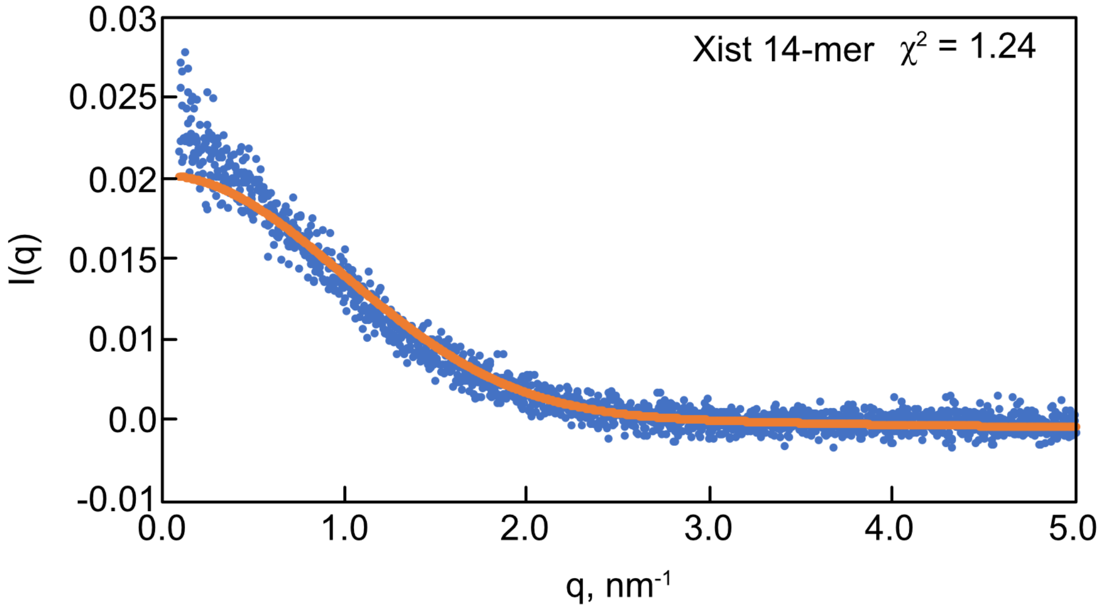


**Supplementary Figure 1.**

Structural model of the Xist A-repeats. CRYSOL goodness of fit curve (orange line) for the 14-mer SAXS curve (blue dots) with model 1 of the NMR resolved 14-mer structure.

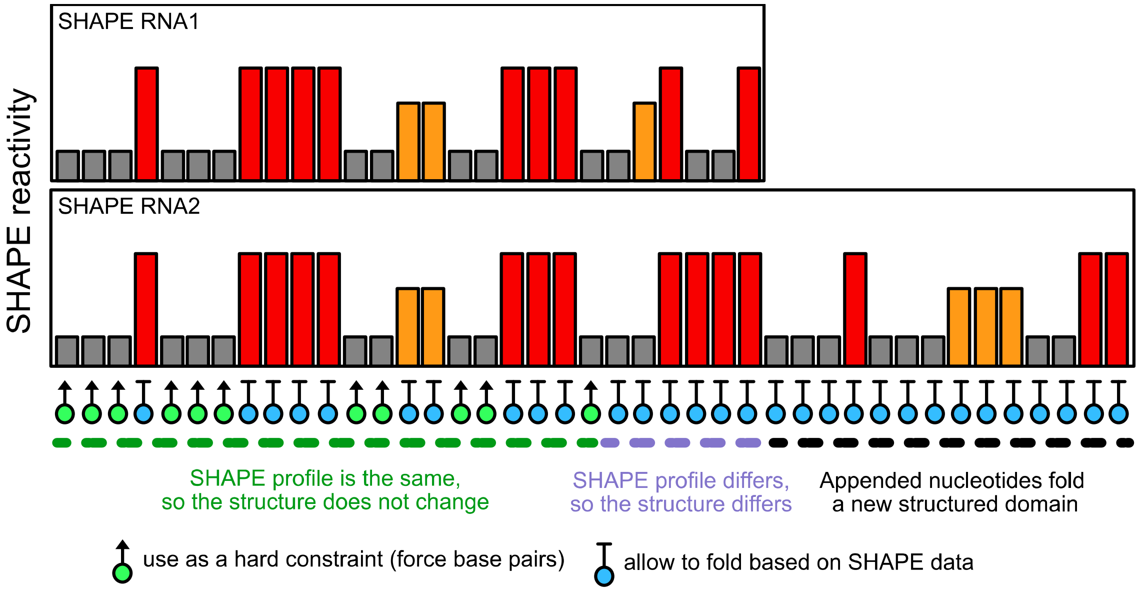


**Supplementary Figure 2.**

Graphical illustration of the approach used to generate the folding restraints to outline the folding of the Xist A-repeats. Structured elements corresponding to clusters at least three nucleotides that maintain a constant SHAPE profiles are held constant in subsequent structure predictions.

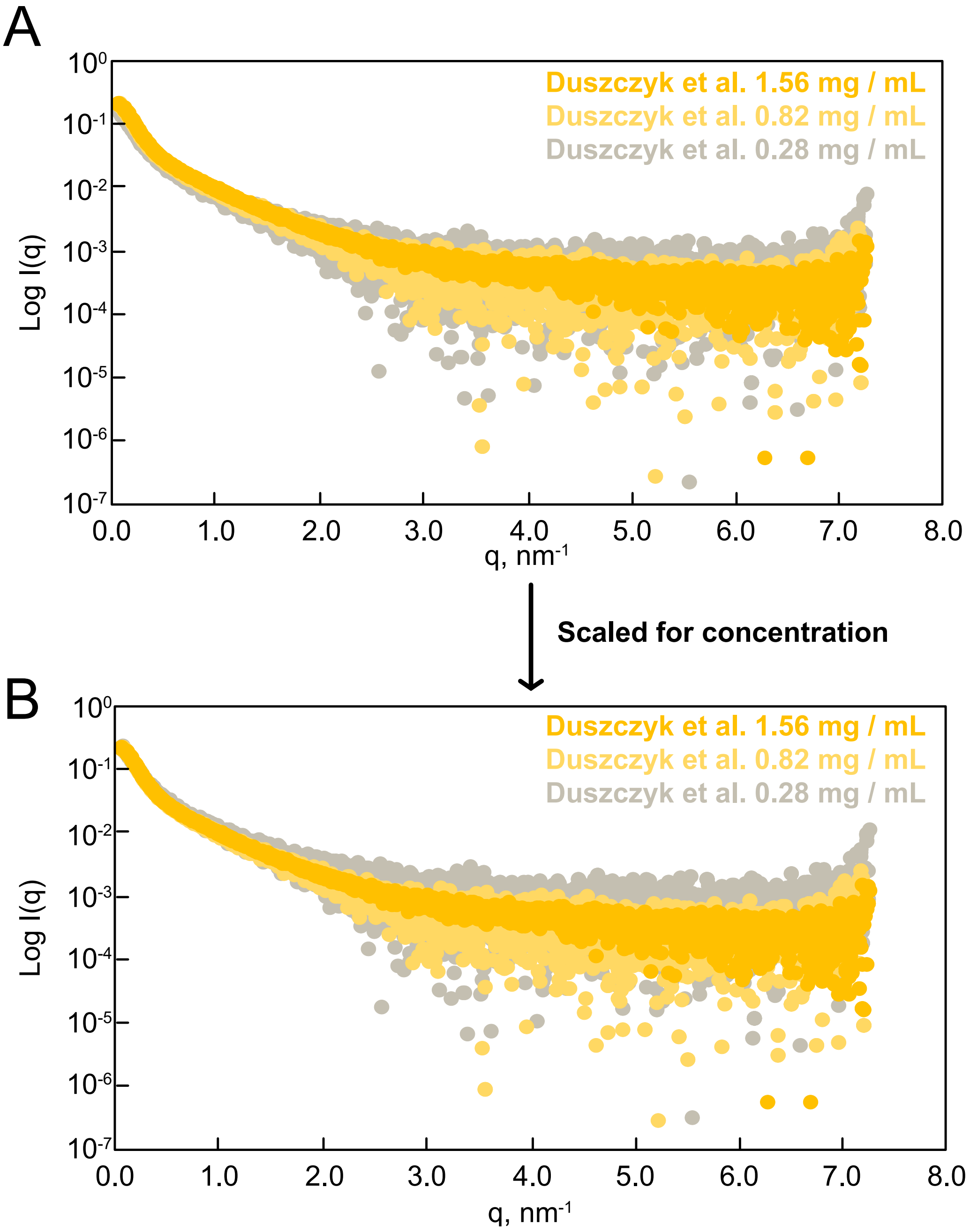


**Supplementary Figure 3.**

Representative concentration series SAXS curves for the Duszczyk et al. buffer conditions of the Xist A-repeat lncRNA at 298K. **(A)** Superposition of the low, medium, and high concentration series data prior to scaling. **(B)** The SAXS curves scaled for concentration.

**Supplementary Table 1.**
**Concentrations of Xist A-repeat arrested RNAs evaluated by SAXS.**

| RNA construct (298 K) | SAXS concentrations (mg/mL) |
|---|---|
| 14-mer | 0.23, 0.36, 0.62 |
| 26-mer | 0.73, 1.12, 1.98 |
| A-rep1 | 0.56, 1.33, 1.85 |
| A-rep2 | 0.47, 0.81, 1.40 |
| A-rep3 | 0.25, 0.45 |
| A-rep4 | 0.48, 0.71, 1.08 |
| A-rep5 | 0.28, 0.48 |
| A-rep6 | 0.44, 0.75, 1.36 |
| A-rep8 | 0.54, 0.89, 1.64 |

**Supplementary Table 2.**
**Comparison of SHAPE reactivity profiles between two consecutive arrested transcripts.**

| Arrested RNA constructs | Pearson R |
|---|---|
| A-rep2: A-rep3 | 0.67 |
| A-rep3: A-rep4 | 0.47 |
| A-rep4: A-rep5 | 0.52 |
| A-rep5: A-rep6 | 0.40 |
| A-rep6: A-rep7 | 0.25 |
| A-rep7: A-rep8 | 0.54 |

**Supplementary Table 3.**
**Concentrations of Xist A-repeat RNA evaluated by SAXS**

| Buffer condition (298 K) | SAXS concentrations (mg/mL) |
|---|---|
| Liu et al., | 0.31, 0.72, 1.37 |
| Dusczyzk et al., | 0.28, 0.82, 1.56 |
| Smola et al., | 0.20, 0.89, 1.55 |
| Maenner et al., | 0.23, 0.71, 2.06 |

**Data S1. (separate file)**

Summary of chemical probing and enzymatic cleavage data for the Xist A-repeat lncRNA.
